# *In Silico* Improvement of Highly Protective Anti-Malarial Antibodies

**DOI:** 10.1101/2022.04.08.487687

**Authors:** Mateo Reveiz, Prabhanshu Tripathi, Lais Da Silva Pereira, Patience Kiyuka, Tracy Liu, Baoshan Zhang, Yongping Yang, Brian G. Bonilla, Marlon Dillon, Myungjin Lee, Chen-Hsiang Shen, Arne Schön, Sven Kratochvil, Facundo D. Batista, Azza H. Idris, Robert A. Seder, Peter D. Kwong, Reda Rawi

## Abstract

Antibody CIS43 binds *Plasmodium falciparum* circumsporozoite protein (PfCSP) and protects against malaria, as recently demonstrated clinically. To improve the efficacy of CIS43, we developed an *in silico* pipeline to optimize the interaction energy of CIS43 to its junctional epitope (peptide 21: PfCSP residues 101-115). Starting from two improved CIS43 variants, recently elicited from a CIS43-germline knock-in mice, single and double amino acid substitutions in the peptide 21-proximal heavy (VH) and light (VL) variable regions were introduced. CIS43-variants, selected on the basis of improved *in silico* interface and stability energies, showed increased affinity to peptide 21 and superior malaria-protective efficacy. The best designed variant, antibody P3-43, was significantly more protective than its template antibody m43.151, with greater liver-burden protection than the current best-in-class (antibody iGL-CIS43.D3). Crystal structures of improved antibodies revealed atomic-level interactions explaining gains in binding affinity. The reported pipeline provides a powerful *in silico* approach to improve antibody functionality.

## INTRODUCTION

Malaria is an infectious disease caused by several species of *Plasmodium* parasites. In 2020, there were 241 million cases of malaria and 627,000 deaths worldwide, with children under 5 accounting for an estimated 80% of all malaria-related deaths in the African region (World Health Organization, 2021). *P. falciparum* is the most prevalent, accounting for ∼50% of cases, and is often resistant to anti-malarial drugs (Phillips et al., 2017). Infection occurs when an infected female *Anopheles* mosquito bites a human and injects sporozoites, which rapidly migrate to the bloodstream. These quickly invade hepatocyte cells in the liver, where they reproduce for 2-10 days before releasing thousands of merozoites to the bloodstream, starting the erythrocytic stage (Cowman et al., 2016).

The *P. falciparum* circumsporozoite protein (PfCSP) is the predominant surface antigen of sporozoites (Plassmeyer et al., 2009), making it a prominent target of vaccines such as RTS,S, which displays the C-terminal portion of PfCSP comprising a number of NANP repeats as well as the C-terminal region (Laurens, 2020; RTSS Clinical Trials Partnership, 2012). It is also the target for multiple monoclonal antibodies such as L9, CIS43 and MGG4 (elicited from the attenuated PfSPZ vaccine), 311 and 317 (elicited from the RTS,S vaccine) and 580g or 663 (from natural exposure) (Kisalu et al., 2018; Oyen et al., 2017; Tan et al., 2018; Triller et al., 2017; Wang et al., 2020). In particular, CIS43 targets the junctional region referred to as peptide 21 (residues 101-115 of PfCSP). The Fc-modified variant, CIS43LS showed promising results in phase I human clinical trials, with a single administration protecting against malaria infection for over 9 months (Gaudinski et al., 2021; Kisalu et al., 2021). More recently (Kratochvil et al., 2021), two variants m42.127 and m43.151 were elicited from a CIS43-germline knock-in mice vaccinated with a KLH-based prototype malaria vaccine candidate. These antibodies were used to design the best-in-class antibody, iGL-CIS43.D3, which was ∼10 fold better than wild-type CIS43 (Kratochvil et al., 2021). These results highlight the potential of CIS43 variants in the fight against malaria. However, the production and distribution of monoclonal antibodies remains challenging (Hooft Van Huijsduijnen et al., 2020). Increased potency of CIS43 variants, however, should allow for lower dosing, reducing production costs and enabling more impactful initiatives to protect vulnerable populations in malaria endemic regions.

Antibody improvement can be performed through multiple methods, both experimental and computational, with increasing computational speeds highlighting the potential of *in silico* approaches. While direct *in silico* simulations of liver burden protection are not yet feasible, it is possible to use interaction energies between *in silico* mutated antibody variants and junctional peptide 21 as a surrogate for binding affinity, which has been shown to correlate with improved liver burden protection (Kratochvil et al., 2021). Because of the large number of potential antibody mutations to be screened, we developed a method that could be scaled for thousands of mutants, through mutating residues and short energy minimizations followed by a single molecular dynamics step to define interface and stability interaction energies. The resulting mutants and their corresponding interaction energies were selected as pareto optimal solutions for experimental antigenic assessment by AlphaLISA and bilayer interferometry (BLI) as well as for functional *in vivo* assessment in a murine malaria challenge model. To further understand the atomic-level interactions leading to higher binding affinity, we performed residue pairwise energy analysis of the resulting *in silico* models, and compared these results experimentally by determining crystal structures of top variant antibodies in complex with peptide 21. Overall, our results demonstrate the ability of the described pipeline to improve antibodies, with the best obtained antibody, P3-43, showing significantly more protection than its template antibody and higher *in vivo* protection than iGL-CIS43.D3, the current best-in-class antibody.

## RESULTS

### *In Silico* Antibody Improvement Pipeline

We developed an *in silico* pipeline to improve the potency of CIS43 antibody variants by optimizing interaction energies to the PfCSP-junctional peptide 21 (**Figure 1A**). Starting with published crystal structures of two template antibodies m42.127 and m43.151 in complex with peptide 21 (PDB ID: 7LKB and 7LKG), we generated all possible single amino acid substitutions in the heavy (VH) and light (VL) variable regions within 12 Å of the peptide (**Figure 1D**). For each mutant we performed a short energy minimization to remove side chain clashes using FoldX and YASARA software (Schymkowitz et al., 2005). Non-bonded van der Waals (vdW) and electrostatic interaction energies between the peptide and the antibody residues within 12 Å of the peptide were calculated (interface metrics). The heavy to heavy, light to light, and heavy to light chain interactions energies were also calculated and treated as stability measures (**Figure 1B, Table 1**). The best single mutants were first selected by taking the top pareto fronts in the 2-dimensional interface energy space. The stability measures in the remaining 6-dimensional space were used as ε-constraints, that is, mutants with larger stability energies than one standard deviation from the average energy were discarded. For the m42.127 template, 42 mutants were found in the top five pareto fronts of which 19 satisfied the ε-constraints (**Figure 1C left**). For the m43.151 template, 23 mutants were found in the top four pareto fronts while 11 were within the allowed stability regions (**Figure 1C right**). The top single mutants were subsequently used to generate double mutants by following a greedy combinatorial approach. From the 19 m42.127-based single variants, 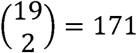 double combinations were assessed and down selected by using the same pareto-based approach described for the single variants. Four final variants of the first pareto front within stability bounds were selected. From the 11 m43.151-based single variants, 55 combinations were assessed for a final of 6 variants (**Figure S1**). Overall, we down-selected a total of 29 single and 6 double mutants (35 total variants) for experimental antigenic assessment (**Figure S1B**).

**Table 1.**
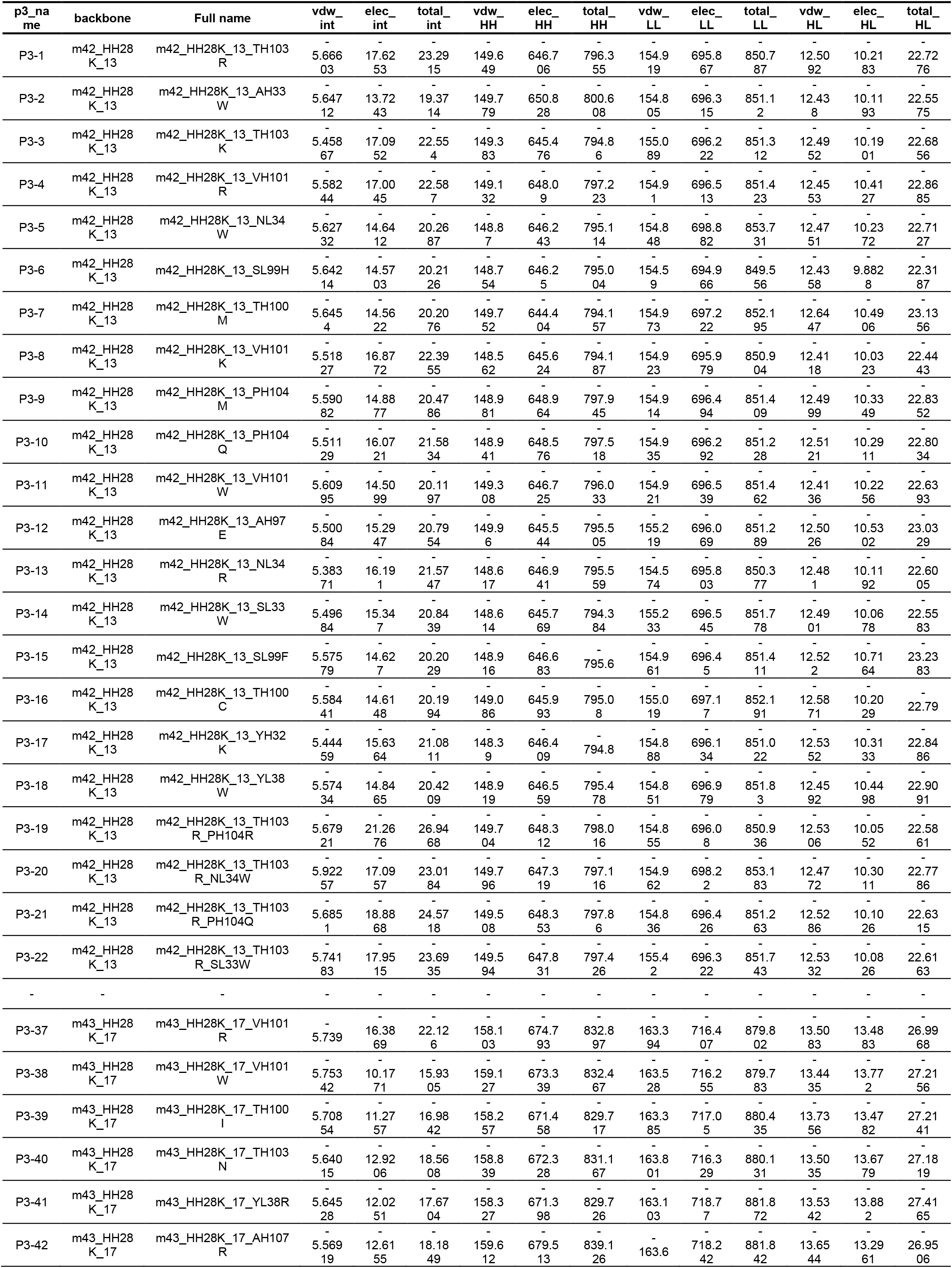
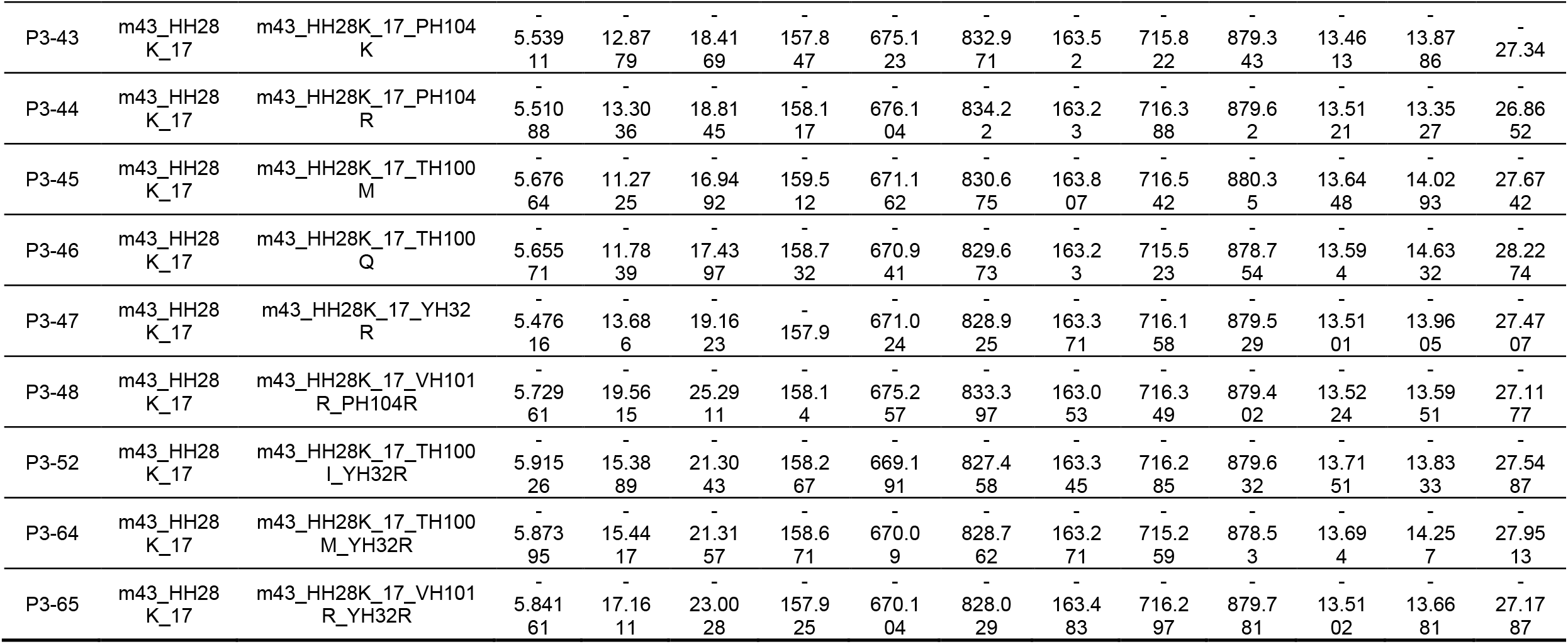
*In silico* energies of selected antibodies.

**Figure 1.**
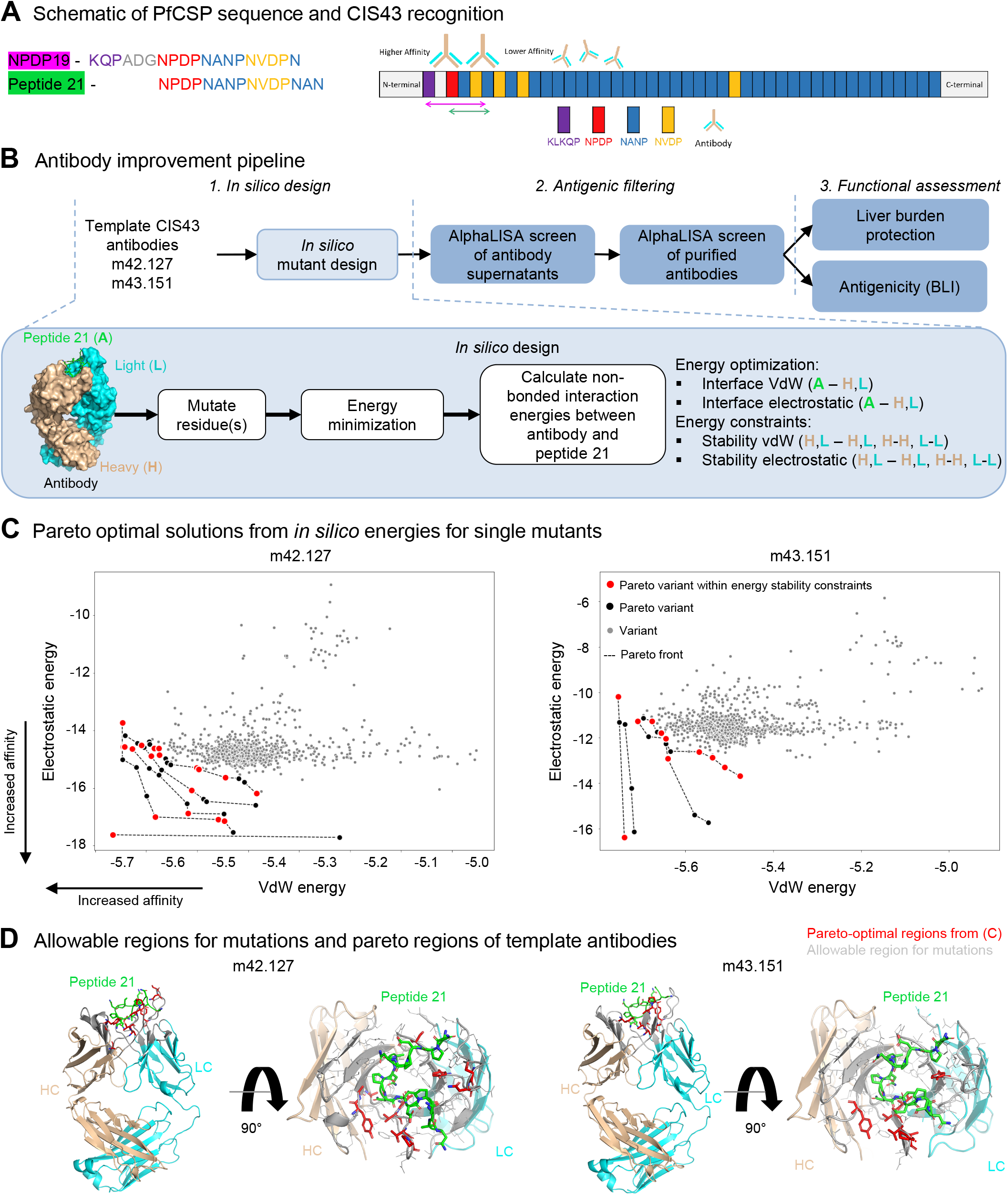
Antibody improvement pipeline identifies CIS43 variants with improved *in silico* non-bonded interaction energies to junctional peptide 21. (A) Schematic of PfCSP sequence and junctional peptides. (B) Antibody improvement pipeline is composed of three sequential steps. First, *in silico* interaction energies are calculated. Second, antigenic filtering down-selects most promising antibody variants. Lastly, functional assessment compares antibody variants against current best-in-class antibodies. (C) Pareto optimal solutions from *in silico* energies for all single mutants based on m42.127 and m43.151 templates. Mutations within the top pareto fronts are depicted in black and red circles. Variants satisfying all stability constraints are highlighted in red. (D) Allowable region for mutations within 12Å of peptide 21 (green) for m42.127 (left) and m43.151 (right) is shown in gray. Residue positions with pareto-optimal energies identified *in silico* are shown in red. See also Figures S1 and S3.

### Antigenic Characterization Confirms Increased Affinity to Junctional Peptide 21

The 35 *in silico* designed antibody variants were produced and antigenically assessed using AlphaLISA, which provided an apparent affinity based on chemiluminescent signals. First, the supernatants of the antibodies were expressed and assessed for their reactivity to the junctional peptide NPDP19, as the affinity to NPDP19 has been shown to have high correlation with function and malaria protection efficacy (Kratochvil et al., 2021). From the initial 35 *in silico* designs, 27 successfully expressed (**Table 2**) and 11 showed improved binding when compared to m43.151 (**Figure S2A**). Subsequently, the 11 top antibody variants were purified and re-assessed against peptide21 using AlphaLISA at two different IgG concentration, in particular 1nm (**Figure S2B**) and 10nm (**Figure S2C**) to avoid hook effects and to allow a more expansive selection of antibodies. Out of the 11 selected antibodies, only antibody P3-44 containing the mutation P100R_H_ did not show improvement over its template antibody m43.151 and was not further considered (**Figure S2B, S2C**). The selected antibodies based on the m42.127 template include five single variants with mutations in the CDRH3 region (T99K_H_, T96M_H_, P100M_H_, P100Q_H_) and one rare mutation in the CDRH1 region (Y32K_H_) (**Figure 2A**). Two multi-mutants were also selected with CDRH3 mutations (T99R_H__P100R_H_ and T99R_H__P100Q_H_). Based on the m43.151 template, four single variants are concentrated in the CDRH3 region (A100cR_H_, P100K_H_, P100R_H_ and T96M_H_). Overall, predictions suggest position 100 in the heavy chain to be important as it was frequently mutated. Most of the mutations identified by the *in silico* pipeline had not been elicited in previous murine models from which the template antibodies where derived (days 13 and 28 from immunization), with the exception of T96M_H_ (**Figure S1B**) (Kratochvil et al., 2021).

**Table 2.**
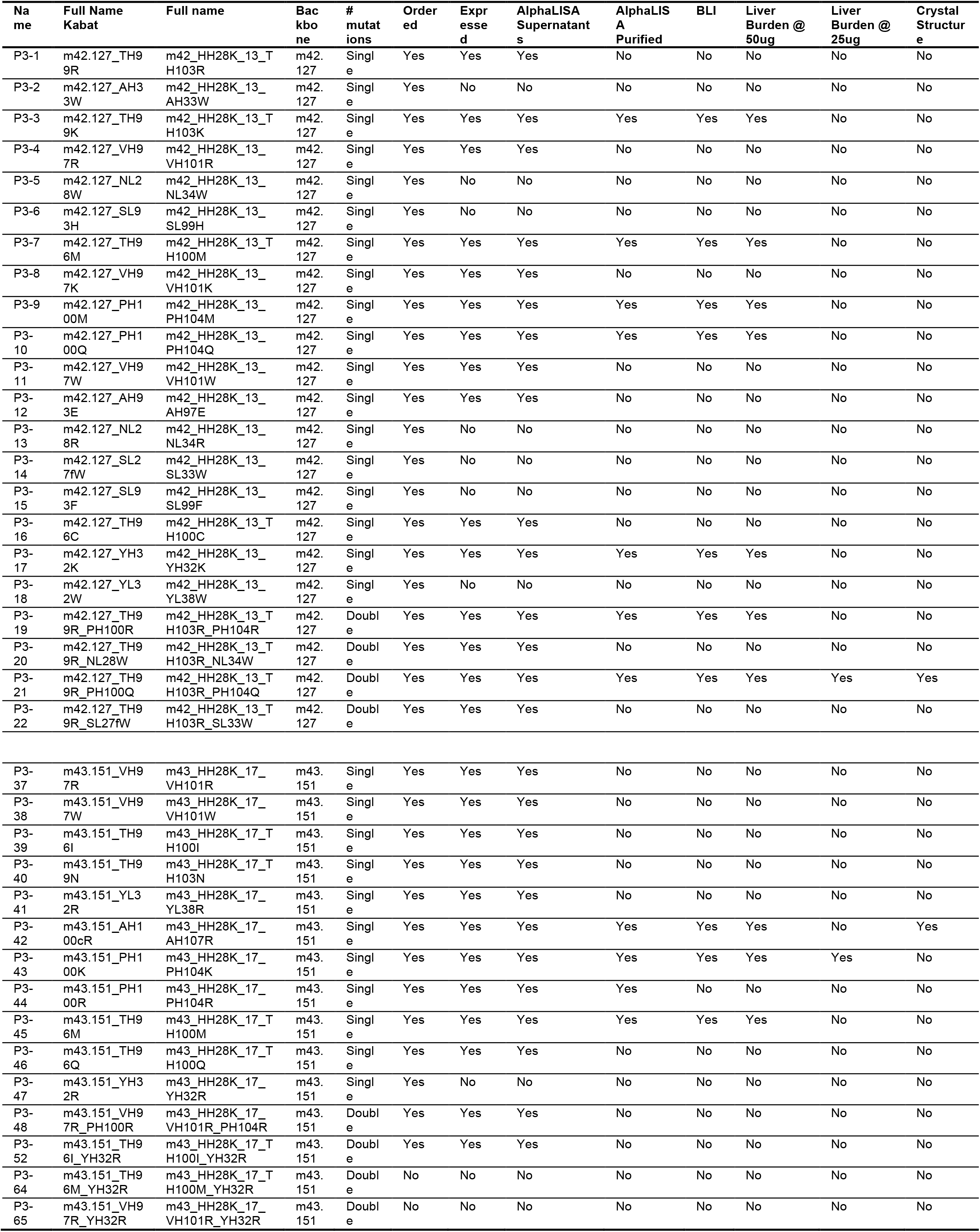
Variants summary and data availability.

**Figure 2.**
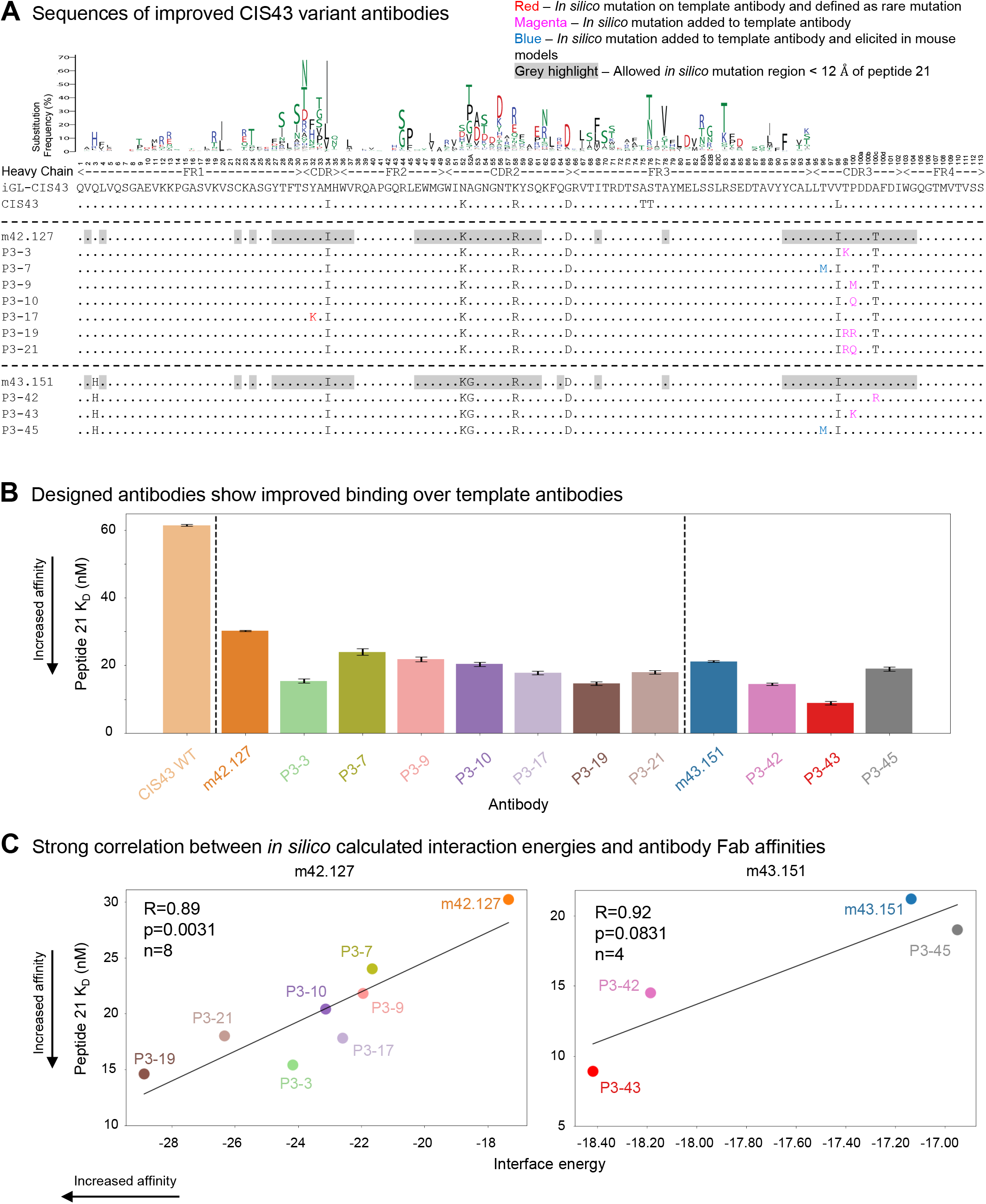
*In silico* designed CIS43 variants show improved binding to peptide 21. (A) Heavy chain sequences for the top mutants as characterized by purified AlphaLISA signals. (B) Biolayer interferometry (BLI) affinity of selected CIS43 variants to peptide 21. Variants are grouped depending on their template antibody. (C) *In silico* interface energy strongly correlates to peptide 21 BLI Kd measurements for m42.127-based variants. See also Figure S2.

Next, we produced the antibody-binding fragments (Fabs) of the antibodies and measured binding affinities to the *in silico* target peptide 21 using bio-layer interferometry (BLI), as well as to NPDP19 and *PfCSPm (****Figure 2B, S3A***). Binding to Peptide 21 was improved for all selected antibodies by ∼2-fold on average with respect to their template antibodies m42.127 (30.2 ± 0.2 nM) and m43.151 (21.2 ± 0.2 nM), with P3-43 showing the tightest binding with 8.9 ± 0.5 nM (**Figure 2B**). Interestingly, the calculated interaction energies showed strong significant correlation with the experimentally determined binding affinities (m42.127: R=0.89, P-value=0.0031; m43.151: R=0.92, P-value=0.0831 with only four measurements) validating the *in silico* approach (**Figure 2C**).

### Functional Characterization of Designed Variants Reveals New Best-In-Class Antibody

As we hypothesized that improved junctional peptide affinity would result in antibodies with higher antimalarial protection capacity, we sought to functionally evaluate the designed variants using an *in vivo* murine challenge model (Wang et al., 2020). Each designed variants with improved BLI affinity were assessed using a group of 5 mice. The template antibodies, m42.127 and m43.151, were used controls, as well as the previous best-in-class antibody iGL-CIS43.D3, naïve (no challenge) and untreated controls. Designed antibodies were intravenously administered 2 hours before malaria challenge. Mice were then challenged with 2000 Pb-PfCSP-GFP/Luc Sporozoites/ IV/ Mouse. Bioluminescence was used to determine malaria liver burden (at day 2) and parasitemia (day 6) (**Figure 3A**).

**Figure 3.**
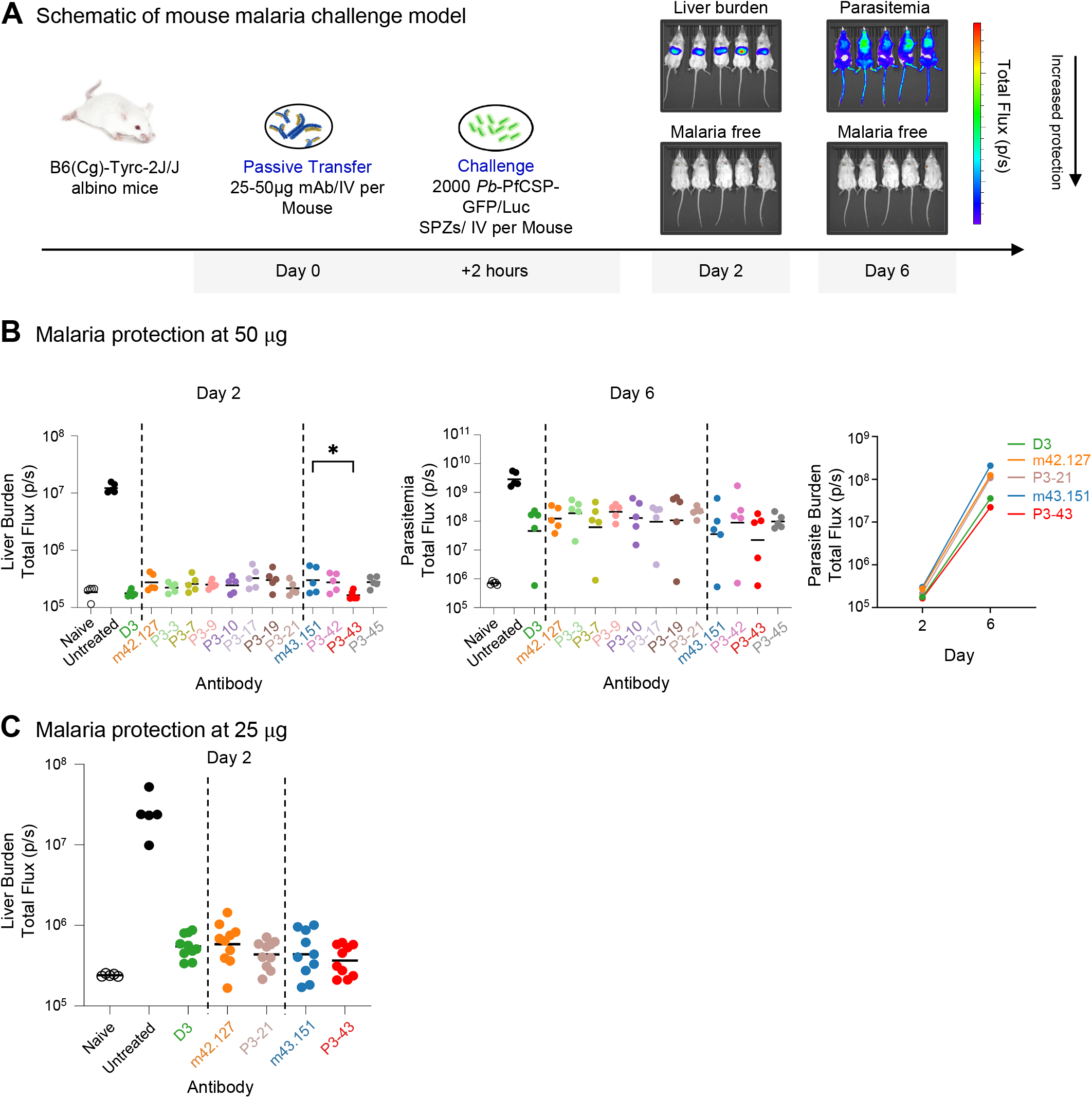
Functional characterization of designed antibodies reveals novel best in class antibody P3-43. (A) Schematic of mouse malaria challenge model. (B) P3-antibodies mediate protection against malaria infection. Mice (n=5/group) were passively infused with 50 *µ*g antibody prior to challenge with transgenic *P. berghei* sporozoites expressing PfCSP and a green fluorescent protein/luciferase fusion protein. Bioluminescence or total flux (photons/second) was quantified at day 2 as a measure of liver stage infection, and again at day 6 to determine parasitemia (blood stage infection). Relative to its template antibody m43.151, P3-43 was statistically superior in liver burden protection assessed using the uncorrected Dunn’s test. (C) Selected P3-antibodies liver burden protection at 25μg dosage. Best P3 antibodies P3-21 and P3-43 were assessed along their template antibodies and best-in-class antibody D3 using a lower concentration of 25 *µ*g. P3-43 is still trending to be the most protective antibody, however, without statistical significance.

Relative to its template antibody m43.151, P3-43 was statistically superior in liver burden protection assessed using the uncorrected Dunn’s test at day 2 for 50µg antibody (**Figure 3B**). The best P3 antibodies for each template, P3-21 and P3-43 were further assessed along their template antibodies and the current best-in-class antibody D3 using a lower concentration of 25µg. P3-43 trended to be the most protective antibody, though without statistical significance (**Figure 3C**).

### *In Silico* Energies and Interatomic Interactions Provide Structural Insight

To understand further the impact of the introduced mutations, we first generated a pairwise interaction energy matrix between peptide 21 and all antibody residues. We then calculated ΔE by subtracting the variant matrix by its corresponding template antibody matrix. For the best m42.127 based variant P3-21 most of the interaction energy gain originates from the introduced residues Arg99_H_ and Gln100_H_, both located in the CDRH3. Their interaction energies account for 63% and 28% for Arg99 and Gln100 respectively (**Figure 4A bottom left**). Interestingly, both Arg99_H_ and Gln100_H_ residues strongly interact with peptide 21 residue Asp11, with Arg99 additionally showing strong interactions to peptide 21 residues Pro8 and Asn9 (**Figure 4A top left**). Regarding the best m43.151 based variant P3-43, the majority of interaction energy gain originates from the introduced residue Lys100_H_ in the CDRH3 with 74% of overall energy contribution (**Figure 4A bottom right**). The energy gain is mostly originating from interactions with peptide residues Asp11 and Ala14 (**Figure 4A top right**). Also, P3-42 shows a similar trend where newly introduced positively charged residue Arg100c_H_ moderately improves the electrostatic energy with negatively charged peptide 21 Asp11 and vdW energy with Asn13 (**Figure S4A top**). In general, the gain in interaction energy originates from electrostatic rather than vdW improvements.

**Figure 4.**
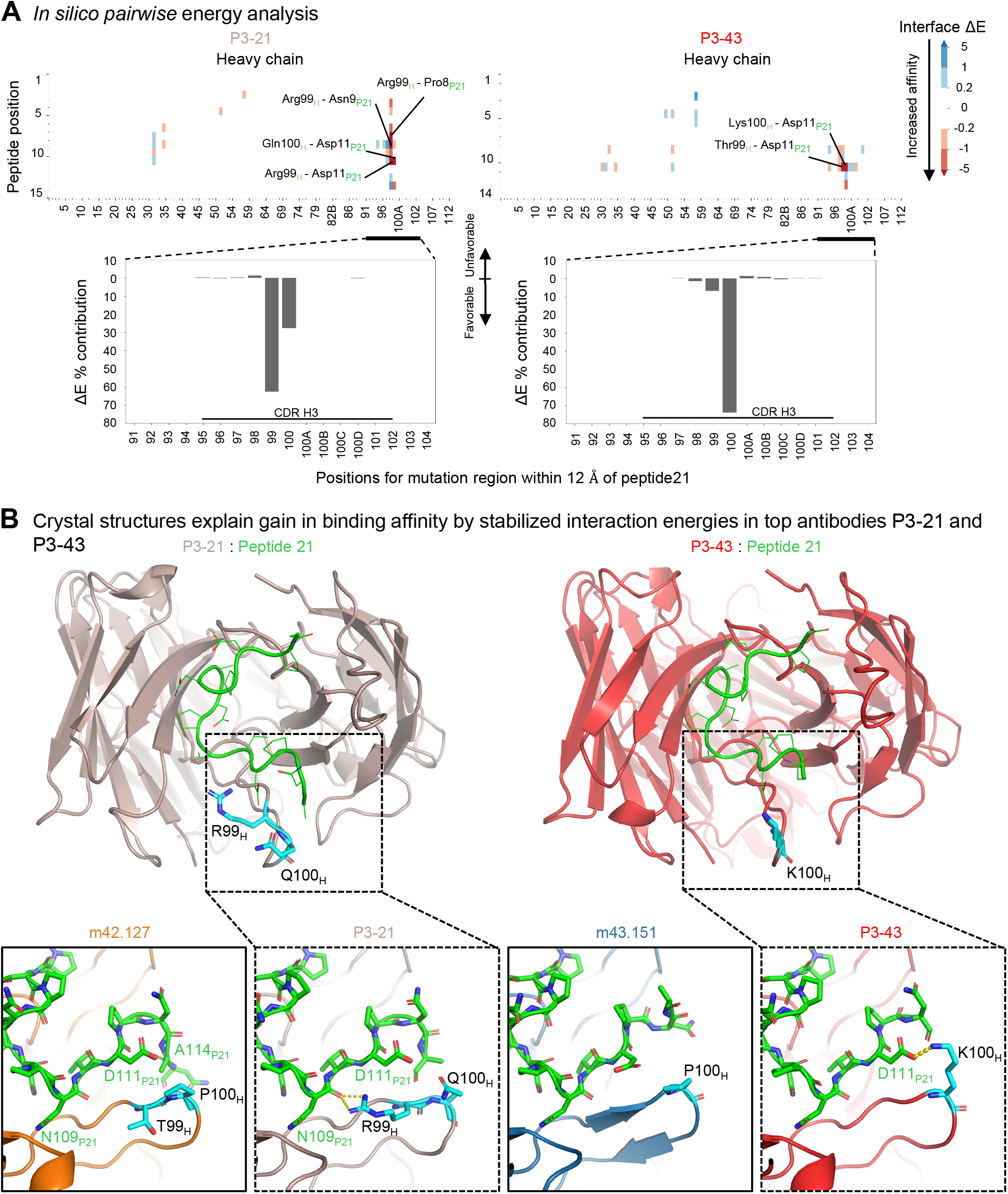
*In silico* energies and crystal structure of P3-21 in complex with peptide 21 depict additional atomic interactions explaining increase in affinity. (A) *In silico pairwise* energy analysis. For each pair of residues between peptide 21 (Y-axis) and the antibody (X-axis), the total interface energy for the antibody is subtracted from the corresponding template antibody. Lower values indicate more favorable interaction energies. In the bar plots below, values are summed across the peptide 21 positions to further examine the CDRH3 region. (B) Crystal structures of P3-21 (top left panel in brown) and P3-43 (top right panel in red) in complex with peptide21 (green) is shown in cartoon illustration. Mutated residues are depicted in cyan stick conformation. Critical binding regions (bottom dotted panels), including heavy chain position 99 and 100 of template antibody m42.127 and P3-21 in complex with peptide21. Amino acid Arg99 in P3-21 introduces new electrostatic interactions with peptide21 backbone atoms. Interatomic interactions from residue 99 in heavy chain and residue 9 in peptide 21 is displayed and compared with that of template m42.127 crystal structure. P3-43 mutation Lys100 introduces new interactions with peptide 21 Asp11. See also Figure S4.

### Structural Analysis Reveal Interactions Leading to Increased Affinity to Junctional Peptide 21

To analyze the atomic-level interactions leading to improved binding, we sought to determine the crystal structures of the top antibodies. We co-crystallized antibody variants P3-21, P3-42, and P3-43 with peptide 21 and determined their structures in complex with peptide 21 to 2.2 Å (**Figure 4B left panel, Table 3)**, 1.8 Å (**Figure S4B)**, and 2.3 Å resolution, respectively (**Figure 4B right panel, Table 3**). The analysis of P3-21 co-complex structure showed additional interactions between the introduced mutations and peptide 21. In particular, the Thr99Arg_H_ mutation allowed the side chain of Arg99_H_ to interact with peptide 21 via hydrogen bonding interactions with the backbone carbonyl of Asn109_P21_ (depicted as yellow dashed lines in **Fig. 4 B left panel**). The introduction of Pro100Gln_H_ mutation is thought to increase flexibility of the CDR-H3 loop, resulting in a weak hydrogen bond between Gln100_H_ and Asp111_P21_. P3-1 variant (m42.127_Thr99Arg_H_) that lacks the Pro100Gln_H_ did not show improved apparent affinity to peptide 21 in the AlphaLISA measurements. P3-43 showed the highest affinity to peptide 21 (Figure 2B) and the structural analysis of P3-43 in complex with peptide 21 revealed a strong salt bridge interaction between the introduced Pro100Lys_H_ mutation and Asp111_P21_, as predicted by the *in silico* model (74% contribution by Lys100_H_ predicted) (**Figure 4B right panel)**. The additional interaction suggested by the *in silico* model between Thr99_H_ and peptide 21 Asp11 was however not visible on the crystal structure as the side chain of Thr99_H_ is facing outwards, while the short energy minimization during the *in silico* MD step led to the Thr99_H_ facing towards the Asp11 residue.

**Table 3:** X-ray crystallography data collection and refinement statistics.

| *PDB accession code* | **P3-21 : Pep21**<br>*7UFN* | **P3-42 : Pep21**<br>*7UFO* | **P3-43 : Pep21**<br>*7UFQ* |
| --- | --- | --- | --- |
| **Data collection** |  |  |  |
| Wavelength | 1 | 1 | 1 |
| Resolution range (Å) | 51.82 - 2.20 (2.28 - 2.20) | 45.52 – 1.8 (1.86-1.80) | 43.52 – 2.30 (2.38 – 2.30) |
| Space group | P 2<sub>1</sub> | C 2<sub>1</sub> | P 2<sub>1</sub> |
| Cell dimensions |  |  |  |
| *a*, *b*, *c* (Å) | 36.7 124.0 103.6 | 94.5 60.7 86.5 | 74.5 61.7 89.6 |
| *a*, *b*, *g* (°) | 90.0, 90.1, 90.0 | 90.0, 106.1, 90.0 | 90, 103.7, 90.0 |
| Unique reflections | 46287 (4700) | 42986 (4118) | 34697 (3381) |
| Multiplicity | 3.6 (3.6) | 3.5 (3.2) | 3.4 (3.6) |
| Completeness (%) | 99.8 (99.9) | 98.2 (94.3) | 97.45 (96.88) |
| *I* / σ*I* | 5.7 (2.0) | 7.5 (1.9) | 8.92 (2.10) |
| Wilson *B*-factor | 25.21 | 15.59 | 32.07 |
| *R*<sub>merge</sub> | 0.159 (0.562) | 0.099 (0.572) | 0.203 (1.017) |
| CC<sub>1/2</sub> | 0.98 (0.79) | 0.99 (0.76) | 0.984 (0.587) |
| **Refinement** |  |  |  |
| Reflections used in refinement | 46921 (4700) | 42944 (4104) | 34476 (3381) |
| Reflections used for R-free | 2448 (248) | 2208 (213) | 1702 (175) |
| *R*<sub>work</sub> | 0.17 (0.13) | 0.19 (0.26) | 0.22 (0.27) |
| *R*<sub>free</sub> | 0.20 (0.17) | 0.22 (0.30) | 0.28 (0.31) |
| Number of non-hydrogen atoms | 6947 | 3811 | 7037 |
| macromolecules | 6891 | 3456 | 6916 |
| ligands | 25 | 22 | 6 |
| solvent | 31 | 333 | 115 |
| Protein residues | 903 | 455 | 906 |
| RMS(bonds) (Å) | 0.009 | 0.007 | 0.010 |
| RMS(angles) (°) | 1.21 | 1.05 | 1.27 |
| Ramachandran favored (%) | 95.29 | 97.30 | 95.41 |
| Ramachandran outliers (%) | 0.11 | 0.22 | 0.0 |
| Average *B*-factor (Å<sup>2</sup>) | 24.74 | 21.92 | 33.70 |
| macromolecules | 24.56 | 21.50 | 33.75 |
| ligands | 61.73 | 26.02 | 20.00 |
| solvent | 35.60 | 25.95 | 31.21 |

For the crystal structure of antibody P3-42, the introduced Arg100c_H_ residue is substantially distant from Asn109_P21_ (**Figure S4B)**. Long-range electrostatic interaction energies may provide a basis for the observed increase in binding affinity, particularly between Asp111_P21_ and Arg100c_H_ as suggested by the *in silico* model.

Overall, the *in silico* model pairwise analysis revealed the atomic-level interactions contributing to the increase in affinity as shown by the corresponding crystal structures.

## DISCUSSION

In this study, we show that an *in silico* energy-based pipeline is able to identify key mutations that improve binding to the junctional peptide 21 as confirmed by antigenic assessment using BLI. Binding improvement was here used as a surrogate function for liver burden protection which does translate to a certain extent as evidenced by antibody variant P3-43 which showed the best peptide 21 binding using BLI and the strongest liver burden protection with ∼10 fold improvement over the antibody WT-CIS43 used in clinical trials. However, liver burden protection cannot be solely explained by improved binding and there are many complex biological factors that can play a role. For example, PfCSP’s extended spiral conformation of the repeat regions allow for complex inter-Fab contacts; mutations to non-antigen positions could enhance or decrease the potency of antibodies (Oyen et al., 2018).

While we were able to improve over the best natural antibodies m42.127 and m42.151, these antibodies were already highly optimized and contained 6 and 7 mutations in the heavy chain respectively, with respect to iGL-CIS43. From a methodological development point of view, it would have been interesting to investigate the performance of the algorithm when starting from germline iGL-CIS43 or WT-CIS43 mature variants for a more direct comparison to the mice models used to discover the template antibodies. However, in this paper we sought to directly optimize the top murine antibodies to rapidly discover new potential clinical products. Nonetheless, this *in silico* approach enables sampling of all potential amino acid substitutions, which is not possible in an *in vivo* system, which is constraint due to other biological factors. Also, our pipeline was set up to efficiently sample many different mutations, with the caveat that we could not perform extensive conformation sampling of the introduced mutants, for instance using molecular dynamics. Hence, we did not account for diverse backbone conformations after introducing large residues, which might affect overall interaction energy contributions.

While this method was not designed to generate *de novo* antibodies and requires an a *priori* binding antibody in complex with its target, we proved that it can be effectively used to refine and optimize antibody variants when used in conjunction with *in vivo* models. This method could also be used in earlier stages to explore and guide the experimental design of animal models as this method has a significant cost and time advantage when compared to other experimental. Interaction energies for one single variant can be calculated in ∼5 minutes using 24 CPUs. Simulating all peptide-proximal mutations using high performance computing settings can be accomplished within days, while smaller computational platforms might require more time. This novel *in silico* pipeline provides a powerful and generally applicable approach to improve antibody functionality and leads in this case to the design of novel antibodies to be used for passive prevention against malaria, by further optimizing CIS43 variants against different epitopes such as peptide 29 or additional antibodies such as L9, but also to additional infectious disease targets.

## ACKNOWLEDGMENTS

We thank J. Stuckey for assistance with figures, and members of the Structural Biology Section and Structural Bioinformatics Core, Vaccine Research Center, for discussions and comments on the manuscript. Support for this work was provided by the Intramural Research Program of the Vaccine Research Center, National Institute of Allergy and Infectious Diseases. This work utilized the computational resources of the NIH HPC Biowulf cluster (http://hpc.nih.gov). Use of sector 22 (Southeast Region Collaborative Access team) at the Advanced Photon Source was supported by the US Department of Energy, Basic Energy Sciences, Office of Science, under contract number W-31-109-Eng-38.

## AUTHOR CONTRIBUTIONS

M.R. and R.R. headed the creation of the *in silico* pipeline, performed computational calculations and led the study. P.T. purified antibodies, performed BLI and AlphaLISA experiments, made Fabs and determined the crystal structures. B.Z. and T.L. performed large scale transfection of antibodies, Y.Y. performed 96-well plate transfections, L.D.S.P., P.K, B.G.B. and M.D. assessed malaria protective efficacy of antibodies in mouse models. M.L. performed the Arpeggio analysis, C.H.S performed sequence analysis, A.S. performed ITC experiments, S.K. and F.D.B. provided template antibodies m42.127 and m43.151. A.H.I. co-led variant PfCSP antibody protective efficacy measurements, R.A.S. oversaw malaria protection studies, P.D.K. and R.R. oversaw the project. M.R. and R.R. provided the main draft of the manuscript, to which all authors provided revisions or comments.

## MATERIALS AND METHODS

### *In silico* Interaction Energy Calculations

Starting with the template pdb files for m42.127 and m42.151 in complex with peptide 21, we first calculated the minimum distance between any antibody residue atom and any peptide atom. Residues within 12 Å were selected and allowed to mutate using the BuildModel command from FoldX software (http://foldxsuite.crg.eu/). The variants were energy minimized using YASARA (http://www.yasara.org/). The reference structures were also modeled and energy minimized by making V1V_H_ and Q1Q_H_ identity mapping mutations. A single molecular dynamics frame was run using the NAMD force field. The energies computed cover both electrostatic and vdW interface energies *F*(*x*) (heavy/light to peptide pairs) and stability energies *G*(*x*) (heavy to heavy, light to light and heavy to light pairs). Energies from both capped C and N termini were ignored to avoid artifacts and more closely match their native state in the PfCSP protein. After computing the interaction energies for all single mutants *x*, the problem becomes to down select the best performing variants by minimizing the multi-objective function *F*(*x*) while constraining the stability function *G*(*x*). To do this, the pareto fronts for the two interface energies were calculated using custom python scripts. A pareto front here is defined as the set of all pareto optimal solutions and in this context, a mutant *u* is a member of the pareto front if:

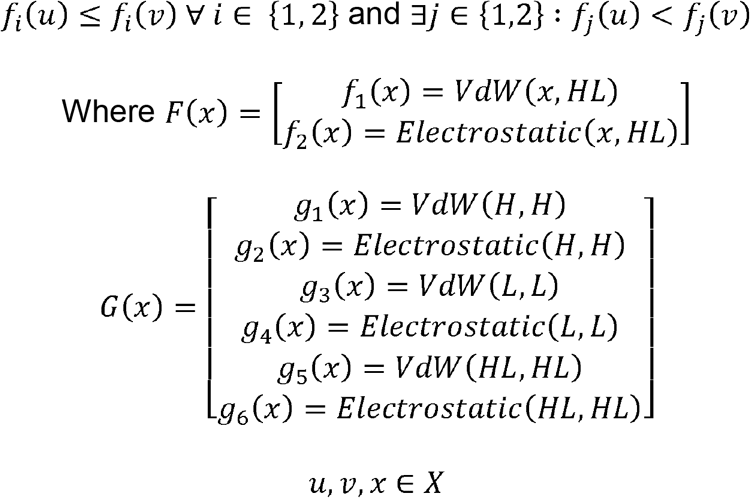

The standard deviation plus the mean of the remaining six stability energies were calculated and used and constraints to the G(u) stability function, such that variant u would be included if *g*_*i*_(*u*) < *µ*_*i*_ + *σ* _*i*_. Here, *µ*_*i*_ is the average of *g*_*i*_ for all variants and *σ*_*i*_ is the standard deviation of *g*_*i*_ for all variants.

Variants that satisfied both the pareto optimality constraints and stability constraints were combined to generate double variants. In other words, we optimize the following multi-objective function:

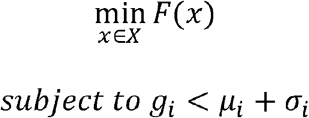

The same pareto optimization and stability constraint method was then used to down select the final double variants.

### Visualization of Intermolecular Interactions in Protein Structures

A web server, Arpeggio (Jubb et al., 2017) was used to visualize the interactions between peptides and antibody heavy/light chains in P3-21 and P3-42.

### Antibody Expression and Purification

Antibody variable heavy chain and light chain sequences were codon optimized, synthesized and cloned into a VRC8400 (CMV/R expression vector)-based IgG1-LS vector as previously described (Kong et al., 2019).The variants were expressed by transient transfection in Expi293 cells (ThermoFisher Scientific, Waltham, MA) using Turbo293 transfection reagent (SPEED BioSystems, Gaithersburg, MD) according to the manufacturer’s recommendation. 50 μg plasmid encoding heavy-chain and 50 μg plasmid encoding light-chain variant genes were mixed with the transfection reagents, added to 100 ml of cells at 2.5 × 10^6^/ml, and incubated in a shaker incubator at 120 rpm, 37°C, 9% CO_2_. At 5 days post-transfection, cell culture supernatant was harvested and purified with a Protein A (GE Healthcare, Chicago, IL) column. The antibody was eluted using IgG Elution Buffer (ThermoFisher Scientific, Waltham, MA) and were brought to neutral pH with 1 M Tris-HCl, pH 8.0. Eluted antibodies were dialyzed against 1xPBS overnight.

### AlphaLISA Characterization of CIS43 Variants

AlphaLISA® (Perkin-Elmer, Waltham, MA) is a bead-based proximity assay in which singlet oxygen molecules, generated by high energy irradiation of Donor beads, transfer to Acceptor beads, which are within a distance of approximately 200 nm. It is a sensitive high throughput screening assay that does not require washing steps. A cascading series of chemical reactions results in a chemiluminescent signal. The antibody variants expressed in 96-well plates were first quantified by biolayer interferometry and the supernatants were subsequently diluted to 6 nM in transfection media. 5 μL of the IgG supernatants were transferred to an OptiPlate-384 assay plate (white opaque, PerkinElmer, Waltham, MA), mixed with 10μL (10 nM final conc.) of biotinylated peptide probe and 10 uL (10 µg/mL final conc.) of Anti-human IgG (Fc specific; Perkin-Elmer, Waltham, MA) acceptor beads. After an hour of incubation at RT, non-shaking, 25 uL (40 µg/mL final conc.) of streptavidin donor beads (Perkin-Elmer, Waltham, MA) were added. The plate was then incubated for 30 min at RT in the dark before the AlphaLISA signal was detected using a SpectraMax® i3x multi-mode microplate reader (Molecular Devices, San Jose, CA). The antibodies selected from the initial screening of supernatants were expressed and purified in large scale. Purified antibodies were diluted to 10 nM in AlphaLISA® buffer (PBS + 0.05% Tween-20 + 0.5 mg/mL BSA) and the AlphaLISA apparent affinity was measured as above.

### Crystallization and Structural Analysis

Antibody Fab and peptide 21 (PfCSP residues 101–115) complexes were prepared by mixing 1:2 molar ratio to a concentration of 15 mg/ml. Crystallization conditions were screened in Hampton Research screening kits, Wizard screening kits, Precipitant Synergy screening kits using a mosquito robot. Crystals initially observed from the wells were manually reproduced. The P3-21: P21 complex crystal grew in 5% isopropanol, 2 M lithium sulfate, 0.1 M magnesium sulfate hydrate and 0.1 M sodium acetate trihydrate pH 4.5; the P3-42: P21 complex crystal grew in 0.1 M TRIS hydrochloride pH 8.5, 2.8 M ammonium sulfate, and P3-43:P21 complex crystal grew in 0.1 M TRIS hydrochloride pH 8.5 and 2.25 M ammonium hydrogen phosphate. Crystals were cryoprotected in 20% glycerol and flash-frozen in liquid nitrogen. Data were collected at a temperature of 100 K and a wavelength of 1.00 Å at the SER-CAT beamline ID-22 (Advanced Photon Source, Argonne National Laboratory, Lemont, IL). Diffraction data were processed with the HKL2000 suite. Structure solution was obtained by molecular replacement with Phaser using CIS43 Fab structures (PDB ID: 6B5M) as a search model. Model building was carried out with Coot. Refinement was carried out with Phenix. Ramachandran statistical analysis indicated that the final structures contained no disallowed residues or no more than 0.22% disallowed residues. Data collection and refinement statistics are shown in **Table 3**.

### Affinity measurements by BLI

Antibody Fab binding affinity to various ligands were measured using biolayer interferometry on an Octet Red384 instrument (fortéBio) with streptavidin capture biosensors (fortéBio) in solid black tilt-well 96-well plates (Geiger Bio-One). Assays were performed with agitation at 30°C. Immobilization of biotinylated PfCSPm, NPDP19, and Pep21, was performed for 60s, followed by a 60s baseline in buffer (PBS + 1% BSA). Association with Fab (serially diluted from 1000 to 62.5 nM) was done for 60s, followed by a dissociation step in buffer for 180s. In all Octet measurements, parallel correction to subtract systematic baseline drift was carried out by subtracting the measurements recorded for a loaded sensor incubated in PBS. Data analysis was carried out using Octet software, version 9.0. Experimental data were fitted globally with a 1:1 Langmuir model of binding for all the antigens except PfCSPm which was fitted with a 2:1 Langmuir model of binding.

### Malaria Challenge Model

After in silico improvement, the mouse malaria challenge model was used to assess (or test) the the protective capacity of the (high affinity) CIS43 variants. The neutralizing capacity of the CIS43 variants against the SPZ were evaluated using 6 to 8-week-old female B6(Cg)-Tyrc-2J/J albino mice (The Jackson Laboratory) as previously described (Wang, et al Immunity). Briefly, mice were treated intravenously via tail vein with 25ug or 50ug mAb diluted in sterile filtered 1X PBS (pH 7.4; total volume 200 ml/mouse). Two hours after antibody infusion, mice were challenged with intravenous injection of 2,000 freshly harvested Pb-PfCSP-GFP/Luc-SPZ into the tail vein.

The malaria burden of infection was measured at 40-42 hours after challenge to determine extent of liver infection, and at 6 days to assess blood stage infection or parasitemia. Each mouse received 150 mL of D-Luciferin (30 mg/mL) intraperitoneally. Ten minutes after the Luciferin injection, mice were imaged under isoflurane anesthesia, using IVIS® Spectrum in vivo imaging system (PerkinElmer). Liver burden and parasitemia were quantified by total flux (photons/sec) expressed by Pb-PfCSP-GFP/Luc-SPZ using the manufacturer’s software (Living Image 4.5, PerkinElmer).

