## Supplemental Figures for "*In Silico* Improvement of Highly Protective Anti-Malarial Antibodies"

**A** Pareto optimal solutions from *in silico* energies for double mutants

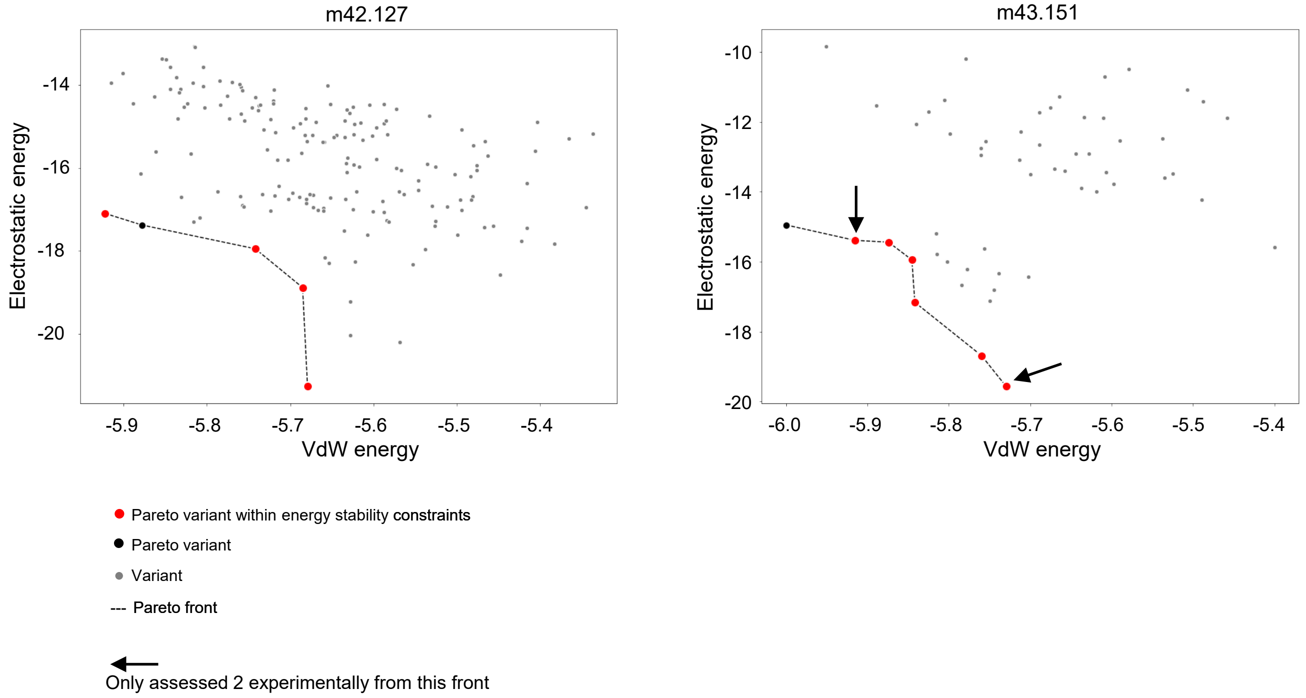


**B** Heavy chain mutations from previous murine models and *in silico methods*

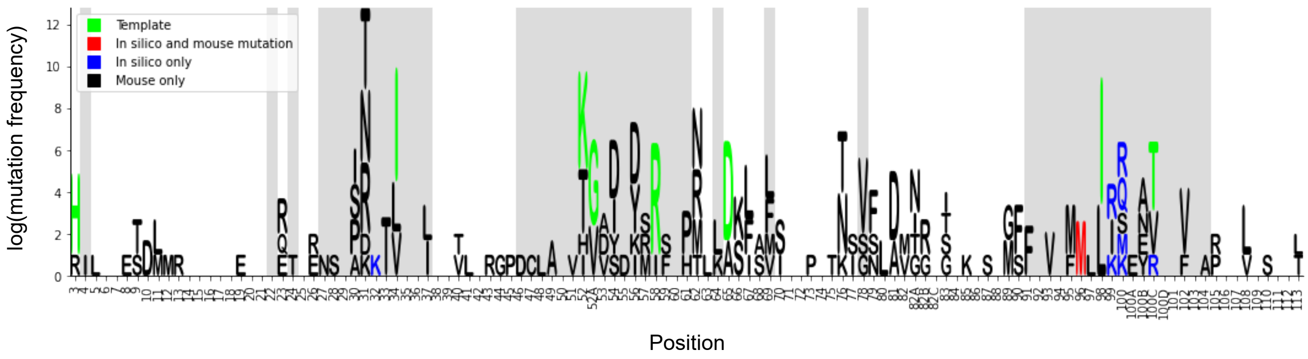


**Figure S1. *In silico* antibody improvement pipeline, related to Figure 1**
(A) Pareto optimal solutions from *in silico* energies for double mutants based on m42.127 (left) and m43.151 (right) templates, related to Figure 1.
(B) Mutation frequency and source of mutation for the heavy chain. Black mutations from previous *in vivo* mouse models. Mouse model mutations contained in the template antibodies m42.127 and m43.151 in green. Mutations identified by the *in silico* pipeline in blue. Mutations identified by both the *in silico* pipeline and mouse models in red. The height of the residues corresponds to the log of the number of antibodies containing the corresponding mutation.

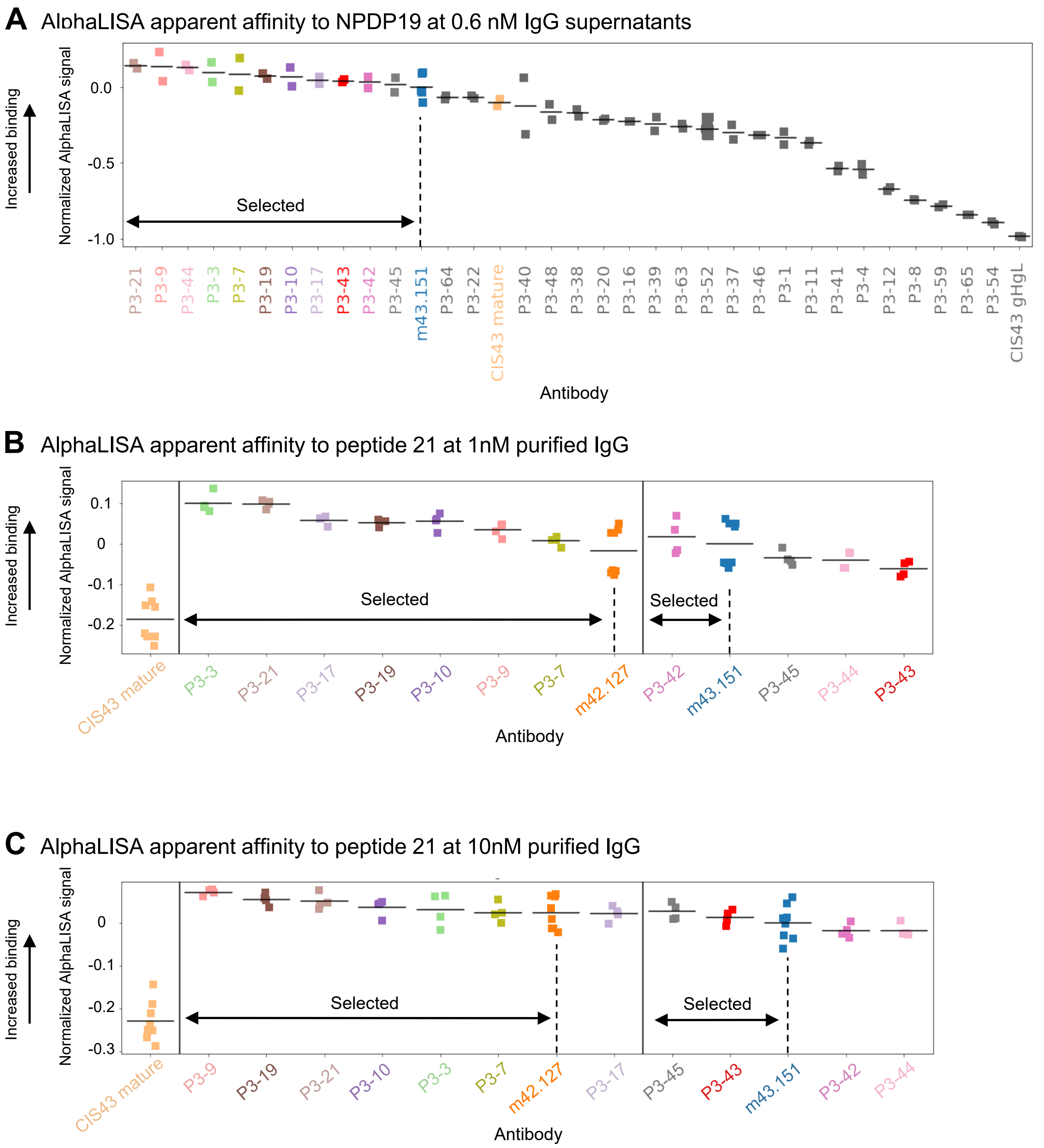


**Figure S2. Antigenic screening using AlphaLISA effectively down-selects best antibodies, related to Figure 2**
(A) AlphaLISA apparent affinity to NPDP19 measured at 0.6 nM with IgG supernatants.
(B) AlphaLISA apparent affinity to Peptide 21 measured at 1 nM with purified IgGs.
(C) AlphaLISA apparent affinity to Peptide 21 measured at 10 nM with purified IgGs.

**A** BLI affinity

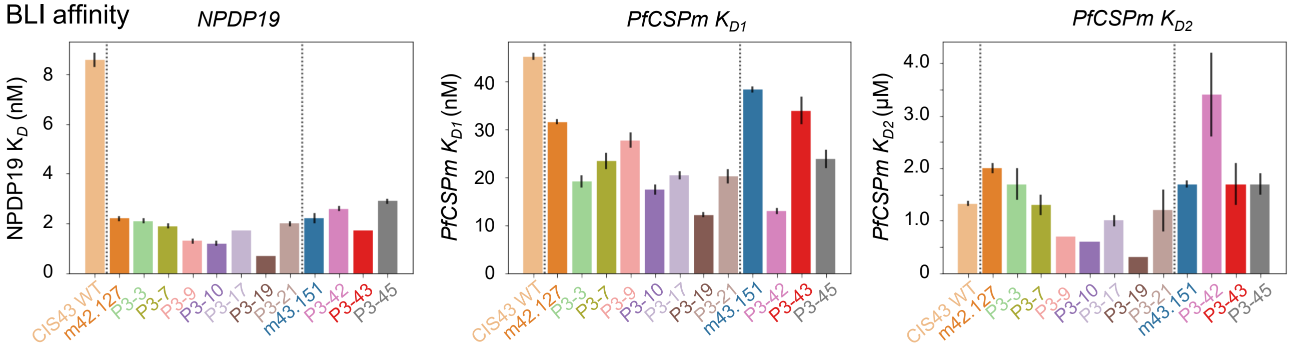


| *Antibody* | *Peptide 21 (nM)* | *NPDP19 (nM)* | *PfCSPm $K_{D1}$ (nM)* | *PfCSPm $K_{D2}$ (μM)* |
|---|---|---|---|---|
| *CIS43 mature** | 61.4 ± 0.3 | 8.6 ± 0.3 | 45.2 ± 0.7 | 1.3 ± 0.0 |
| *CIS43 gHgL** | 635.0 ± 48.7 | 27.2 ± 140 | 679.0 ± 73 | 5.0 ± 0.5 |
| *m42.127** | 30.2 ± 0.2 | 2.2 ± 0.1 | 31.6 ± 0.6 | *m42.127** |
| *P3-3* | 15.4 ± 0.6 | 2.1 ± 0.1 | 19.1 ± 1.3 | 1.7 ± 0.3 |
| *P3-7* | 24.0 ± 1.0 | 1.9 ± 0.1 | 23.4 ± 1.6 | 1.3 ± 0.2 |
| *P3-9* | 21.8 ± 0.7 | 1.3 ± 0.1 | 27.7 ± 1.6 | 0.7 ± 0.0 |
| *P3-10* | 20.4 ± 0.6 | 1.2 ± 0.1 | 17.4 ± 1.0 | 0.6 ± 0.0 |
| *P3-17* | 17.8 ± 0.5 | 1.7 ± 0.0 | 20.4 ± 0.8 | 1.0 ± 0.1 |
| *P3-19* | 14.6 ± 0.5 | 0.7 ± 0.0 | 12.2 ± 0.6 | 0.3 ± 0.0 |
| *P3-21* | 18.0 ± 0.5 | 2.0 ± 0.1 | 20.2 ± 1.5 | 1.2 ± 0.4 |
| *m43.151** | 21.2 ± 0.2 | 2.2 ± 0.2 | 38.3 ± 0.7 | 1.7 ± 0.066 |
| *P3-42* | 14.5 ± 0.3 | 2.6 ± 0.1 | 13.0 ± 0.6 | 3.4 ± 0.8 |
| *P3-43* | 8.9 ± 0.5 | 1.7 ± 0.0 | 33.9 ± 2.9 | 1.7 ± 0.4 |
| *P3-45* | 19.0 ± 0.6 | 2.9 ± 0.1 | 23.9 ± 1.9 | 1.7 ± 0.2 |

* The affinity data for these antibodies is from Kratochvil et al, 2021, *Immunity*.

**B** Isothermal titration calorimetry (ITC)

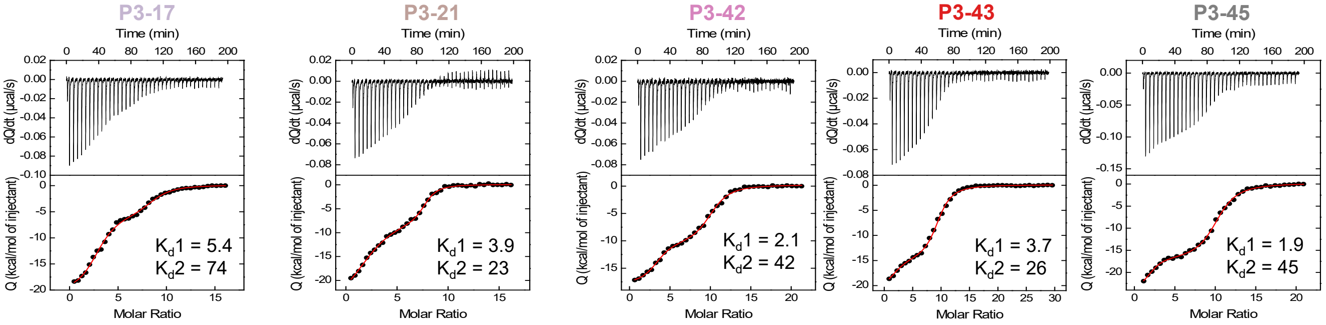


| Antibody | Kd1 (nM) | ΔG1 (kcal/mol) | ΔH1 (kcal/mol) | -TΔS1 (kcal/mol) | N1 | Kd2 (nM) | ΔG2 (kcal/mol) | ΔH2 (kcal/mol) | -TΔS2 (kcal/mol) | N2 |
|---|---|---|---|---|---|---|---|---|---|---|
| *P3-17* | 5.4 | -11.3 | -21.1 | +9.8 | 2.8 | 74 | -9.7 | -5.8 | -3.9 | 5.7 |
| *P3-21* | 3.9 | -11.5 | -26.4 | +14.9 | 2.2 | 23 | -10.4 | -7.2 | -3.2 | 5.4 |
| *P3-42* | 2.1 | -11.8 | -18.4 | +6.6 | 3.3 | 42 | -10.1 | -10.7 | +0.6 | 6.7 |
| *P3-43* | 3.7 | -11.5 | -22.4 | +10.9 | 1.9 | 26 | -10.4 | -14.7 | +4.3 | 7.4 |
| *P3-45* | 1.9 | -11.9 | -24.5 | +12.6 | 1.9 | 45 | -10.0 | -17.3 | +7.3 | 8.2 |

**Figure S3. Biolayer interferometry (BLI) affinity and Isothermal titration calorimetry (ITC) of selected CIS43 Antibodies to junctional peptides and *PfCSP*, related to Figure 1**
(A) Bio-layer interferometry (BLI) affinity of improved CIS43 variants was determined against NPDP19 and *PfCSPm*.
(B) Isothermal titration calorimetry of CSP_SA_mut with various improved CIS43 variants. The affinities and stoichiometry are shown for both KD1 and KD2 in nM.

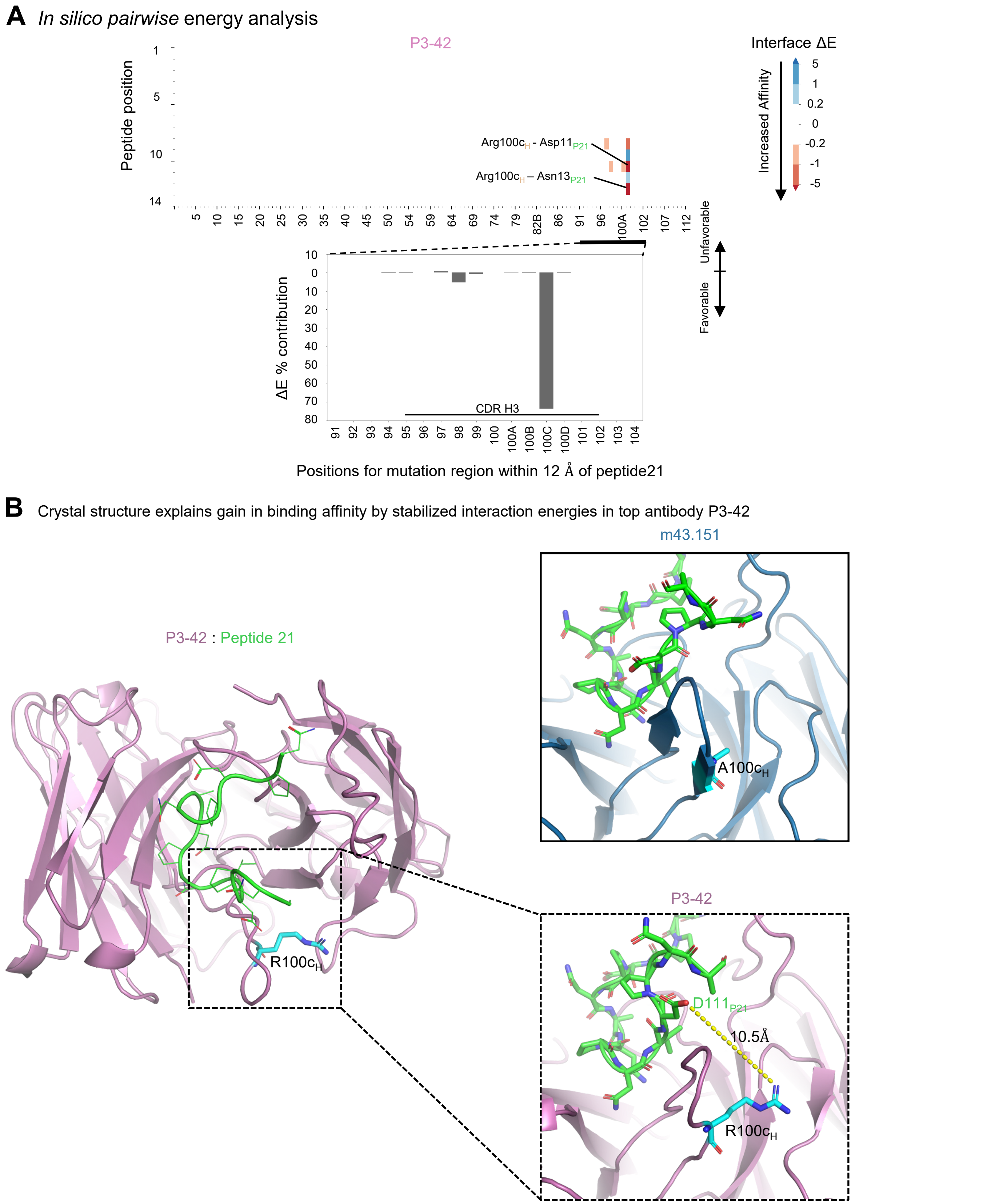


**Figure S4. *In silico* energies and crystal structure of P3-42 in complex with peptide 21 depict additional atomic interactions potentially explaining increase in affinity, related to Figure 4.**

(A) *In silico pairwise* energy analysis. For each pair of residues between peptide 21 (Y-axis) and the antibody (X-axis), the total interface energy for the antibody is subtracted from the corresponding template antibody. Lower values indicate more favorable interaction energies. In the bar plots below, values are summed across the peptide 21 positions to further examine the CDRH3 region.

(B) Crystal structures of P3-42 and its template antibody m43.151 are displayed in cartoon illustration. Interatomic interactions from residue 100c in heavy chain and residue 11 in peptide 21 is displayed (bottom right) and compared with that of template m42.151 crystal structure (top right).
